# Human gut bifidobacteria inhibit the growth of the opportunistic fungal pathogen *Candida albicans*

**DOI:** 10.1101/2021.12.21.473717

**Authors:** Liviana Ricci, Joanna Mackie, Gillian E. Donachie, Ambre Chapuis, Kristýna Mezerová, Megan D. Lenardon, Alistair J. P. Brown, Sylvia H. Duncan, Alan W. Walker

## Abstract

The human gut microbiota protects the host from invading pathogens and the overgrowth of indigenous opportunistic species via mechanisms such as competition for nutrients and by production of antimicrobial compounds. Here, we investigated the antagonist activity of human gut bacteria towards *Candida albicans*, an opportunistic fungal pathogen that can cause severe infections and mortality in susceptible patients. Co-culture batch incubations of *C. albicans* in the presence of faecal microbiota from six different healthy individuals revealed varying levels of inhibitory activity against *C. albicans*. 16S rRNA gene sequence profiling of these faecal co-culture bacterial communities showed that the *Bifidobacteriaceae* family, and *Bifidobacterium adolescentis* in particular, were most correlated with antagonistic activity against *C. albicans*. Follow up mechanistic studies confirmed that culture supernatants of *Bifidobacterium* species, particularly *B. adolescentis*, inhibited *C. albicans in vitro* under both aerobic and anaerobic conditions. Production of the fermentation acids acetate and lactate, together with the concomitant decrease in pH, were strong drivers of the inhibitory activity. Bifidobacteria may therefore represent attractive targets for the development of probiotics and prebiotic interventions tailored to enhance inhibitory activity against *C. albicans in vivo*.

## INTRODUCTION

The human colon harbours a diverse microbiota that is dominated by obligate anaerobic bacteria (Whitman, Coleman and Wiebe 1998; Pasolli *et al*. 2019). The main energy sources for these gut microbes are non-digestible carbohydrates that resist digestion in the small intestine and become available for bacterial fermentation in the proximal colon (Flint *et al*. 2015). These substrates are fermented by the gut microbiota to produce short-chain fatty acids (SCFAs), such as acetate, propionate and butyrate, and other fermentation acids such as lactate (Cummings 1981). SCFAs provide the host with up to 5–10% of their total daily energy requirement (Mortensen and Clausen 1996), and positively impact intestinal and systemic host health (Cummings 1981; Koh *et al*. 2016).

The intestinal microbiota also contributes to host health by bolstering resistance against colonisation of the gut by exogenous pathogens (Bohnhoff, Miller and Martin 1964; Buffie *et al*. 2015). This phenomenon, termed colonisation resistance, can prevent pathogens from establishing and replicating in the gut, or from reaching the densities required to invade deeper tissues and cause overt disease (Bohnhoff, Miller and Martin 1964). Colonisation resistance is multifactorial, involving mechanisms such as the direct production of antimicrobial compounds (Rea *et al*. 2010; Donia and Fischbach 2015), competition for adhesion receptors on the gut epithelium (Ventura *et al*. 2016), and direct competition for niches and nutrients required for the growth of competing pathogenic bacteria (Freter *et al*. 1983; Wilson and Perini 1988; Deriu *et al*. 2013; Maltby *et al*. 2013). Additional mechanisms of colonisation resistance include the creation of a less favourable gut environment, for example lowering the luminal pH through the production of SCFAs (Cherrington *et al*. 1991; Roe *et al*. 2002; Rivera-Chávez *et al*. 2016), or depleting free molecular oxygen, which can prevent the overgrowth and virulence gene expression of some pathogenic microbes (Marteyn *et al*. 2011; Rivera-Chávez *et al*. 2016). Furthermore, human gut commensals are instrumental in the training and modulation of the host immune system (Kau *et al*. 2011; Thaiss *et al*. 2016), inducing the release of host antimicrobial compounds (Cash *et al*. 2006; Fan *et al*. 2015), and in stimulating epithelial barrier reinforcement and repair (Rossi *et al*. 2015; Geirnaert *et al*. 2017). Importantly, microbiota-mediated colonisation resistance can be weakened by various environmental factors and insults, such as Western-style diet (Martinez-Medina *et al*. 2014), antibiotic therapy (Bohnhoff, Miller and Martin 1964; Vollaard EJ, Clasener HA 1992), and acute and chronic inflammatory conditions (Stecher *et al*. 2007; Carroll *et al*. 2012).

*Candida albicans* is a diploid polymorphic fungus and a common opportunistic pathogen of humans, with an estimated annual incidence of 700,000 cases of *Candida* bloodstream infections globally (Guinea 2014). In susceptible patient cohorts, including premature infants and those undergoing chemo-or immune-therapy, organ or stem cell transplants, or abdominal surgery or trauma, *C. albicans* infections can be particularly devastating, with mortality rates of 46–75% following systemic spread, even with antifungal drug interventions (Brown *et al*. 2012). The incidence of *C. albicans* infections has increased in vulnerable subjects over the past few decades (Low and Rotstein 2011) alongside the emergence of other clinically important *Candida* spp., such as *C. auris* (Pfaller *et al*. 2000; Heaney *et al*. 2020). Furthermore, a significant increase of isolates with resistance to common antifungal agents has been observed (Whaley *et al*. 2016).

Despite the pathogenic potential of *C. albicans*, it exists harmlessly in the gastrointestinal tract (GIT) of 40–80% of healthy individuals in Western countries, predominantly in the yeast form, and with cell counts that do not typically exceed 10^4–5^ colony forming units (CFU)/g faeces (Odds *et al*. 1989; Mason *et al*. 2012; Neville, d’Enfert and Bougnoux 2015; Harnett, Myers and Rolfe 2017; Nash *et al*. 2017). The GIT is therefore a natural reservoir of *C. albicans* (Hube 2004; Odds 2010) but, in health, its overgrowth is suppressed by the gut microbiota via colonisation resistance (Kennedy and Volz 1985a; Fan *et al*. 2015). However, conditions such as weakened immunity, increased permeability of the intestinal mucosal barrier, and/or perturbation of microbiota-mediated colonisation resistance via receipt of broad spectrum antibiotics can favour *C. albicans* pathogenesis (Samonis *et al*. 1994; León *et al*. 2009; Gammelsrud *et al*. 2011; d’Enfert *et al*. 2020). Furthermore, systemic candidiasis is often reported to derive from a preceding expansion of *Candida* spp. in the GIT and subsequent translocation from the intestinal niche into the bloodstream (Miranda *et al*. 2009; Zhai *et al*. 2020). GIT colonisation by *C. albicans* is therefore a major risk factor for systemic candidiasis (Pittet *et al*. 1994).

Given the importance of the intestinal niche as a reservoir for systemic dissemination, and the known suppressive effects of the indigenous microbiota on the colonisation of the gut by *C. albicans* in health (Fan *et al*. 2015), we here assessed the potential of the human gut microbiota, and individual gut anaerobe species, to suppress the growth of this opportunistic pathogen *in vitro*. We identified specific bacterial isolates, including *Bifidobacterium adolescentis*, in faecal samples of healthy individuals that inhibit *C. albicans* growth *in vitro*, and revealed the involvement of gut bacterial fermentation acids and pH in this process. These findings suggest that it may be possible to enhance colonisation resistance against *C. albicans* invasive infection using targeted probiotics and/or dietary modulation of endogenous species with antagonistic activity against this opportunistic fungal pathogen.

## MATERIALS AND METHODS

### Ethics

Faecal sample collections used for isolation of human gut anaerobes, and for co-culture experiments with *C. albicans* were approved by the Ethical Review Panel of the Rowett Institute under study number 5946. All donors were received no antibiotic treatment for at least 6 months prior to faecal donation.

### Cultivation of *C. albicans* strain SC5314

*C. albicans* strain SC5314 (Gillum, Tsay and Kirsch 1984) was prepared by plating 2-10 µl of frozen glycerol stock on YPD plates (1% w/v yeast extract (Oxoid LP0021, Basingstoke, UK), 2% w/v mycological peptone (Oxoid LP0040), 2% w/v D-glucose, and 2% w/v agar No. 2 (Oxoid LP0012)) and incubating at 30°C for 48 h. A single colony was transferred from the Petri dish into NGY broth (0.1% yeast extract (Oxoid LP0021), 0.1% neopeptone (Difco, Franklin Lakes, NJ, USA), and 0.4% w/v D-glucose) (MacCallum *et al*. 2006) and incubated at 30°C, with shaking at 200 rpm, overnight. The concentration of *C. albicans* cells in suspension (cells/ml) was estimated by counting using a haemocytometer. Yeast growth was assessed by measuring optical density of the cultures at a wavelength of 600 nm using a spectrophotometer. For determination of *C. albicans* CFUs in samples, cells were plated on Sabouraud dextrose agar (SDA) (4% (w/v) D-glucose, 1% (w/v) mycological peptone, 2% (w/v) agar No. 2, pH 5.6).

### Batch co-cultures of *C. albicans* and mixed faecal microbiota from healthy donors

Co-cultures of *C. albicans* and mixed faecal microbiota were performed in duplicate for each faecal donor in anaerobically sealed Wheaton bottles containing complex anaerobic medium. The medium contained (amounts given are for 1 L): oat spelt xylan (0.6 g; Sigma-Aldrich, St. Louis, MO, USA), pectin (citrus, 0.6 g; Sigma-Aldrich), amylopectin (0.6 g; Sigma-Aldrich), arabinogalactan (larch, 0.6 g; Sigma-Aldrich), potato starch (5.0 g; Sigma-Aldrich), inulin (0.6 g; Sigma-Aldrich), porcine mucin (0.5 g; Sigma-Aldrich), casein hydrolysate (0.5 g; Fluka, Charlotte, NC, USA), peptone water (0.5 g; Oxoid), K_2_HPO_4_ (2.0 g; BDH, Dubai, UAE), NaHCO_3_ (0.2 g; Sigma-Aldrich), NaCl (4.5 g; Fisher Scientific), MgSO_4_ • 7H_2_O (0.5 g; BDH), CaCl_2_ • 2H_2_O (0.45 g; Sigma-Aldrich), FeSO_4_ • 7H_2_O (0.005 g; Hopkin & Willams, UK), haemin (0.01 g; Sigma-Aldrich), bile salts (0.05 g, Oxoid), 0.1% w/v resazurin (0.6 ml), antifoam A (Y-30, 0.5 ml; Sigma-Aldrich), and dH_2_O to 1 L. The pH was adjusted to 6.5 (using 1 mM HCl and 1 mM NaOH, as appropriate) before dispensing the medium anaerobically and autoclaving. After autoclaving, the medium was supplemented with 2 mL mineral solution (150 mg EDTA, 60 mg FeSO_4_ • 7H_2_O, 3.0 mg ZnSO_4_ • 7H_2_O, 0.9 mg MnCl_2_ • 7H_2_O, 9.0 mg boric acid, 6.0 mg CoCl_2_ • 6H_2_O, 0.3 mg CuCl_2_ • 2H_2_O, 0.6 mg NiCl_2_ • 6H_2_O, 0.9 mg NaMoO_4_ • 2H_2_O, and dH_2_O to 300 mL), 1.4 mL vitamin solution (0.2 g menadione, 0.4 g biotin, 0.4 g pantothenate, 2.0 g nicotinamide, 0.1 g vitamin B_12_, 0.8 g thiamine, 1.0 g *p*-aminobenzoic acid, and dH_2_O to 200 mL), and additional components (2 µg folic acid, 2000 µg inositol, 400 µg niacin, 400 µg pyridoxine HCl, 200 µg riboflavin, 100 µg potassium iodide, and 200 µg ferric chloride). In addition, each Wheaton bottle was supplemented with 40 mL filter-sterilised reducing solution to ensure anaerobic conditions (0.5 g cysteine, 3.0 g NaHCO_3_, and dH_2_O to 40 mL).

*C. albicans* cells from an over-night culture grown in YPD broth were washed in sterile PBS, counted using a haemocytometer, and inoculated into 50 ml anaerobic media in Wheaton bottles at a final concentration of 5 × 10^6^ cells/ml (except for one pilot experiment where the inoculum was 5 × 10^5^ cells/ml, see Results section for more details). Faecal samples were obtained from six different donors. 10% (w/v) faecal slurries were prepared in gentleMACS™ M tubes (Miltenyi Biotech, Auburn, CA, USA) by homogenisation in anaerobic PBS (PBS containing 0.05% cysteine). Faecal homogenates were centrifuged at 500 g for 5 min and the liquid faecal component was injected into the Wheaton bottles using a sterile syringe (to give a 0.02% faecal suspension at baseline). The inoculated Wheaton bottles were incubated at 35°C for 48 h with gentle shaking at 75 rpm. Measurements of *C. albicans* colony forming units (CFUs) were carried out at t=0, 24 and 48 h by plating ten-fold serial dilutions on SDA plates supplemented with 34 µg/ml chloramphenicol. CFUs were counted after aerobic incubation at 30°C for 2-3 d.

### 16S rRNA gene sequencing of co-cultured incubation samples

The faecal inocula from healthy donors used in the co-culture experiments, and from the two biological replicate samples collected after 24 and 48 h of incubation with *C. albicans*, were analysed by Illumina MiSeq-based 16S rRNA gene profiling, targeting the V1–V2 region of the gene. Genomic DNA was extracted using the FastDNA™ SPIN Kit for Soil (MP Biomedicals, Irvine, CA, USA) following the manufacturer’s instructions. Barcoded fusion primers containing adaptors for downstream Illumina MiSeq sequencing MiSeq-27F (5′-AATGATACGGCGACCACCGAGATCTACACTATGGTAATTCCAGMGTTYGATYMTG GCTCAG-3′) and MiSeq-338R (5′-CAAGCAGAAGACGGCATACGAGAT-barcode-AGTCAGTCAGAAGCTGCCTCCCGTAGGAGT-3′) were used for PCR amplification of 16S rRNA genes from extracted DNA. PCR was performed using Q5 Taq polymerase (New England Biolabs, Ipswich, MA), with the following cycling conditions: 98°C for 2 min; followed by 20 cycles at 98°C for 30 s, 50°C for 30 s, and 72°C for 90 s; with a final extension at 72°C for 5 min. Each sample was amplified in quadruplicate; the four reactions were pooled and PCR products were ethanol precipitated to generate a single PCR amplicon tube per sample. The PCR products were then quantified using a Qubit 2.0 fluorometer (Life Technologies, Carlsbad, CA, USA), and a sequencing master mix was prepared by mixing the samples in equimolar amounts, which was then sequenced at the Centre for Genome-Enabled Biology and Medicine (CGEBM) at the University of Aberdeen (Aberdeen, UK). For sequencing, an Illumina MiSeq machine was used, with 2 × 250 bp read length. The raw output sequence data are available from the European Nucleotide Archive, under the project accession number PRJEB48351. Individual sample accession numbers are given in **Table S1_Supplementary Data**.

### Analysis of 16S rRNA gene amplicon data

The raw read data in fastq format were analysed using the open-source software Mothur (Schloss *et al*. 2009). For both of the timepoints after co-culture, the two experimental replicates were pooled into single samples for final analyses as no statistically significant differences were detected between replicates. Briefly, contigs were created using the make.contigs command and low quality contigs (such as with length <280 or >470 bases, containing at least one “N”, and polymeric stretches >7 bases) were filtered out using screen.seqs. The contigs were aligned against the SILVA reference (https://www.arb-silva.de/) (Quast *et al*. 2013), and operational taxonomic units (OTUs) were generated at a 97% similarity cut-off level, with a pre-clustering step of diffs=3 to reduce the impact of sequencing errors. Chimera removal software was not used as abundant OTUs corresponding to bifidobacteria were mistaken for chimeric sequences. Instead, the split.abund command was used to filter out low-abundance sequences that appeared less than 10 times in the dataset. All samples were rarefied to 9171 reads for subsequent comparative analyses. Samples derived from the D1 and D3 faecal inocula samples generated far fewer reads than this, so were excluded from the final analyses. Taxonomic classifications were assigned to each OTU by mapping against the RDP reference database (Cole *et al*. 2014). Taxonomies for selected OTUs were also validated by manually checking representative sequences using BLAST searches against the NCBI nucleotide database (https://blast.ncbi.nlm.nih.gov/Blast.cgi), and the Ribosomal Database Project (Johnson *et al*. 2008; Cole *et al*. 2014). Alpha-diversity measures, and phylotype analyses at the phylum, family and genus levels were carried out using Mothur. The final OTU table, phylum, family, genus and alpha-diversity results for each sample are shown in **Table S1_Supplementary Data**. The faecal and enriched microbial community co-culture samples were assigned to the categories ‘benign’ or ‘antagonistic’ according to the extent of the inhibition shown against *C. albicans*. Putative biomarkers at different taxonomic levels that correlated with antagonistic activity against *C. albicans* were assessed using LEfSe (Segata *et al*. 2011) as implemented in Mothur.

### Culturing of human gut anaerobes

The gut anaerobes tested in the current study included isolates from the Rowett Institute (Aberdeen, UK) strain collection or purchased from DSMZ (Braunschweig, Germany) (**Table S2_Supplementary Data**). The isolates were revived from stocks, anaerobically, in Hungate tubes containing M2GSC medium supplemented with 10% v/v clarified bovine rumen fluid (Bryant 1972; Miyazaki *et al*. 1997). Inoculated cultures were incubated at 37°C in a static 5% CO_2_ incubator overnight (NuAire, Plymouth, MN, USA). Cell growth was monitored by measuring optical density at 650 nm (OD_650_) using a spectrophotometer (Novaspec II, Amersham BioSciences UK Ltd., Little Chalfont, UK).

Some of the anaerobic bacteria tested for anti-*Candida* activity in this study were newly isolated from the stool samples of two consenting adults (D3 and DM1). For each donor, 10-fold serial faecal dilutions were prepared in M2 medium (Hobson 1969) with no added carbon source. Each preparation was then used to inoculate five different agar plates: fastidious anaerobe agar (FAA, LAB M Ltd, Heywood, UK) supplemented with 5% v/v horse blood and 0.5% w/v menadione; FAA supplemented with 5% v/v horse blood; brain heart infusion (BHI, Oxoid); M2GSC (Miyazaki *et al*. 1997); and M2GSC supplemented with 0.5% w/v haemin and 0.5% w/v menadione. The plates were incubated in an anaerobic cabinet (Don Whitley Scientific, Bingley, UK) for 48 h. In parallel, faecal dilutions were pre-incubated in M2-AXOS diluting broth (M2 supplemented with 0.2% w/v arabinoxylan oligosaccharides; Cargill, Wayzata, MN, USA) before streaking. After 4 d of incubation, single colonies were selected and picked onto duplicates agar plates of the same type of culture medium they were first grown on. Half of these duplicate plates were left to grow in the anaerobic cabinet, while the remaining plates were incubated aerobically, at 37°C, for up to 48 h. At the end of the incubation, the growth on anaerobic plates was compared with that on the aerobic counterparts to screen for strictly anaerobic isolates. Single colonies were picked from plates that only showed anaerobic growth and then grown in Hungate tubes containing either M2GSC medium supplemented with 0.5% w/v haemin and 0.5% w/v menadione, fastidious anaerobe broth supplemented with 5% v/v horse blood and 0.5% w/v menadione, or BHI broth. DNA was extracted from the collected cultures using the FastDNA™ SPIN Kit for Soil (MP Biomedicals) and 16S rRNA genes were Sanger sequenced (Eurofins Genomics) for taxonomic identification. Culturing conditions used to obtain each of the novel isolates are shown in **Table S3_Supplementary Data**.

### Inhibition of *C. albicans* growth by gut bacterial supernatants and gut bacterial fermentation acids

In order to assess the effect of individual gut bacterial isolates on the growth of *C. albicans* SC5314, anaerobes of interest (**Table S2_Supplementary Data**) were cultured in tubes with anaerobic M2GSC medium at 37°C overnight. The individual culture supernatants were then collected after centrifugation at 658 × *g* for 10 min. The supernatants were filter-sterilised by passing through 0.2 µm syringe-driven filter units (Millex, Merck Millipore Ltd, Kenilworth, NJ, USA) to remove residual bacterial cells. *C. albicans* cells pre-grown in NGY to an OD_600_ of 0.8-0.95 were diluted 1 in 100 in fresh NGY medium and 100 µL was transferred to wells of 96-well microtitre plates (CoStar, Washington, WA, USA). The *C. albicans* suspensions were incubated with an equal amount of filter-sterilised bacterial culture supernatant, or fresh NGY medium as a control, to assess the fungal growth, with technical replicates. The 96-well plates were incubated anaerobically in a temperature-controlled plate reader at 37°C (Epoch 2 Microplate Spectrophotometer, BioTek, Swindon, UK). For each test and technical replicate, the growth of *C. albicans* was calculated by subtracting the OD_600_ value at time 0 from that measured after 24 h (T24–T0). The percentage growth of the fungus in fresh NGY medium in the absence of bacterial supernatant was set as 100% growth reference for each repeat run, and uninoculated filter-sterilised M2GSC medium was used as a control.

The impact of gut bacterial fermentation acids on *C. albicans* growth was assessed by monitoring fungal growth in the presence of a mixed solution of 45 mM sodium acetate (Sigma-Aldrich), 15 mM lactate (Sigma-Aldrich), and 10 mM sodium formate (VWR BDH Chemicals, Merck), supplemented with 0.4% w/v glucose, in addition to individual acids plus 0.4% w/v glucose. The pH of all solutions or NGY medium was adjusted using 1 M NaOH and 1 M HCl, as appropriate, to 4, 5, 6 or 7, and checked using a pH meter (Denver Instrument, Denver, CO, USA).

### Quantification of fermentation acids in gut bacterial culture supernatants using gas chromatography

The culture supernatants of the tested gut bacterial isolates were analysed by capillary gas chromatography (GC) to quantify the production of fermentation acids. To determine the concentrations of SCFAs and lactate, the samples were first derivatised as described elsewhere (Richardson *et al*. 1989). Briefly, 1 mL of a culture supernatant was placed in a Sorvall screw-capped tube and 50 µL of 0.1 M 2-ethylbutyric acid was added as an internal standard. Concentrations of derivatised fatty acids were determined after a double step extraction of organic acids in 0.5 mL of HCl and 2 mL of diethyl ether per sample, and quantification of their tertiary butyldimethylsisyl (*t*-BDMS) derivatives using capillary GC apparatus (Agilent 6890; Agilent Technologies, Santa Clara). Two technical replicates of an external standard (acetic acid, propionic acid, iso-butyric acid, *n*-butyric acid, iso-valeric acid, *n*-valeric acid, sodium formate, lithium lactate, and sodium succinate) were analysed alongside the samples in each GC run to assess quality of the extraction.

### Statistical analyses

The non-parametric Kruskal–Wallis test, followed by Dunn’s post-hoc test, were used to analyse data from assays on the inhibition of *C. albicans* growth by gut bacterial supernatants, and to compare *C. albicans* growth in the absence and presence of gut anaerobe supernatants, using Prism v8.4.1 (GraphPad, San Diego, CA, USA). To test for associations between percent *C. albicans* growth and the gut bacterial culture supernatants, a Spearman correlation was computed using Prism v8.4.1 (GraphPad). Exact P-values obtained using the Spearman correlation test were corrected using the two-stage linear step-up procedure of Benjamini, Krieger and Yekutieli (false discovery rate approach, with Q=5%). Parameters included the OD of microbial cultures, pH, and fermentation acid levels (acetate, formate, and lactate; separately and combined), as quantified in the culture supernatants using GC.

## RESULTS

### Inhibitory activity of cultivated faecal microbiota on *Candida albicans* growth varies between faecal donors

To establish whether the gut microbiota from different individuals vary in their ability to suppress the growth of *C. albicans*, we performed co-culturing experiments in batch culture, where *C. albicans* SC5314 cells were incubated for up to 48 h alongside faecal inocula from six healthy adults. The co-cultures were performed under anaerobic conditions in a complex growth medium designed to mimic the human colon environment. The viability of *C. albicans* cells was assessed by determining CFUs following plating onto SDA medium plus chloramphenicol at t=0 and after 24 h and 48 h incubation with or without homogenised faecal inocula.

An initial experiment was conducted with a stool sample from a single healthy volunteer (Donor 1). As shown in **Figure 1A**, the co-culture of *C. albicans* (inoculated at 5 × 10^5^ cells/ml) with faecal material from Donor 1 showed a clear reduction in the fungal CFUs after 44 h incubation. However, viable cell counts were also reduced at the end of the control incubation when *C. albicans* was grown alone (**Figure 1**, black lines), albeit the reduction was lower than that observed in co-culture. Subsequent experiments, assessing the impact of faecal inocula from five additional donors, were therefore performed using ten times more *C. albicans* cells (inoculated at 5 × 10^6^ cells/ml), which was sufficient to maintain significant *C. albicans* CFUs throughout the experiments (**Figure 1B**). In the control samples, without the faecal inoculum, *C. albicans* CFUs remained relatively constant throughout the 48 h incubation period, with counts around 2.5 × 10^6^ CFU/mL, indicating that the colon-mimicking growth medium and anaerobic conditions did not kill *C. albicans* (**Figure 1B**, black lines). The experiment also revealed that the faecal microbiota from different individuals affected *C. albicans* viable counts to markedly differing degrees after 44 h of co-culture (**Figure 1B**, orange, red, green, brown and blue lines). The faecal inoculum from Donor 5 resulted in the strongest inhibitory effect on *C. albicans* growth, with a 1000-fold (3-log) reduction of *Candida* CFUs at the end of the incubation period (1 × 10^3^ CFU/mL). Co-cultures with faecal inocula from Donors 3, 4, and 6 also resulted in a decrease in *C. albicans* CFUs (between 4 and 20-fold decrease). In contrast, the faecal inoculum from Donor 2 resulted in no effect on *C. albicans* growth, which was comparable with that of the no faecal inoculum control, suggesting that the gut bacteria cultured from the faecal inoculum of this individual did not impair the fungal survival under the tested conditions. We conclude that the cultivated faecal samples from healthy individuals differed in their ability to inhibit the survival of *C. albicans*.

**Figure 1:**
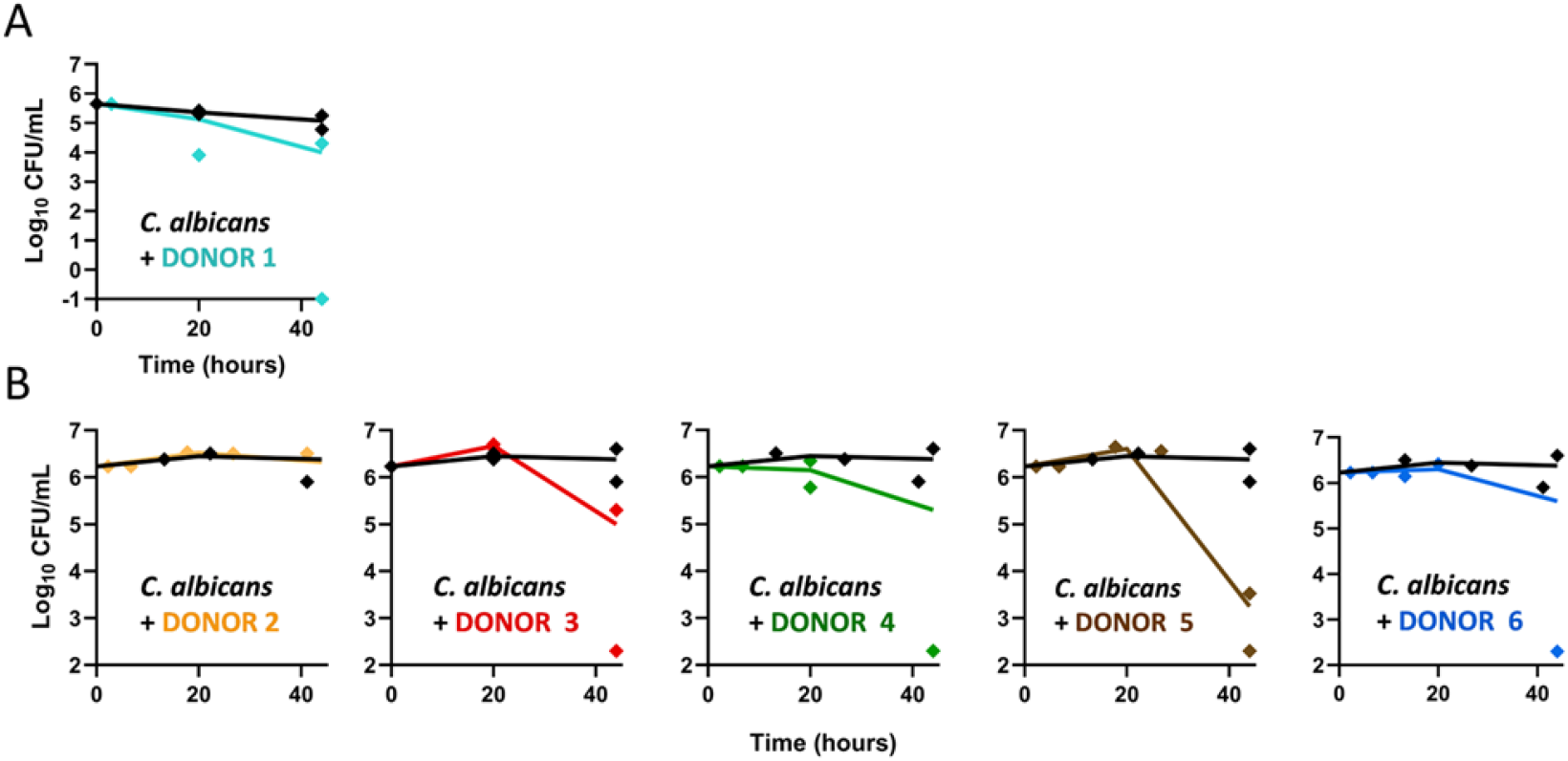
Faecal inocula from healthy donors resulted in varying killing activity against *C. albicans* cells *in vitro*. *C. albicans* was cultured with faecal inocula from six different individuals (Donor 1 – 6), or with no faecal inocula as controls (black lines). Each data point (diamonds) represents *C. albicans* CFU/mL at sampled time points, while the line connects the means at each time point, calculated from two independent CFU measurements. Data were transformed to Log_10_ (y-axis). A) *C. albicans* was inoculated into the anaerobic medium at a density of 5 × 10^5^ cells/ml. B) *C. albicans* was inoculated into the anaerobic medium at a concentration of 5 × 10^6^ cells/ml.

### Variance in faecal microbiota composition may impact colonisation resistance against *C. albicans*

The differing extent of inhibition of *C. albicans* growth observed in co-cultures with faecal inocula from different donors might result from differences in the cultured species composition and, consequently, their metabolic activities. Therefore, we used 16S rRNA gene-based sequence profiling to analyse the bacterial communities present in the initial faecal inocula from the different donors and in the co-culture batch samples after one and two days of incubation. The analysis revealed that, as anticipated, at the OTU level, the initial faecal inoculum samples contained the highest alpha diversity, which then became reduced as certain bacterial taxa were selectively enriched during co-incubation (**Figure 2A**; **Table S1_Supplementary Data**).

**Figure 2:**
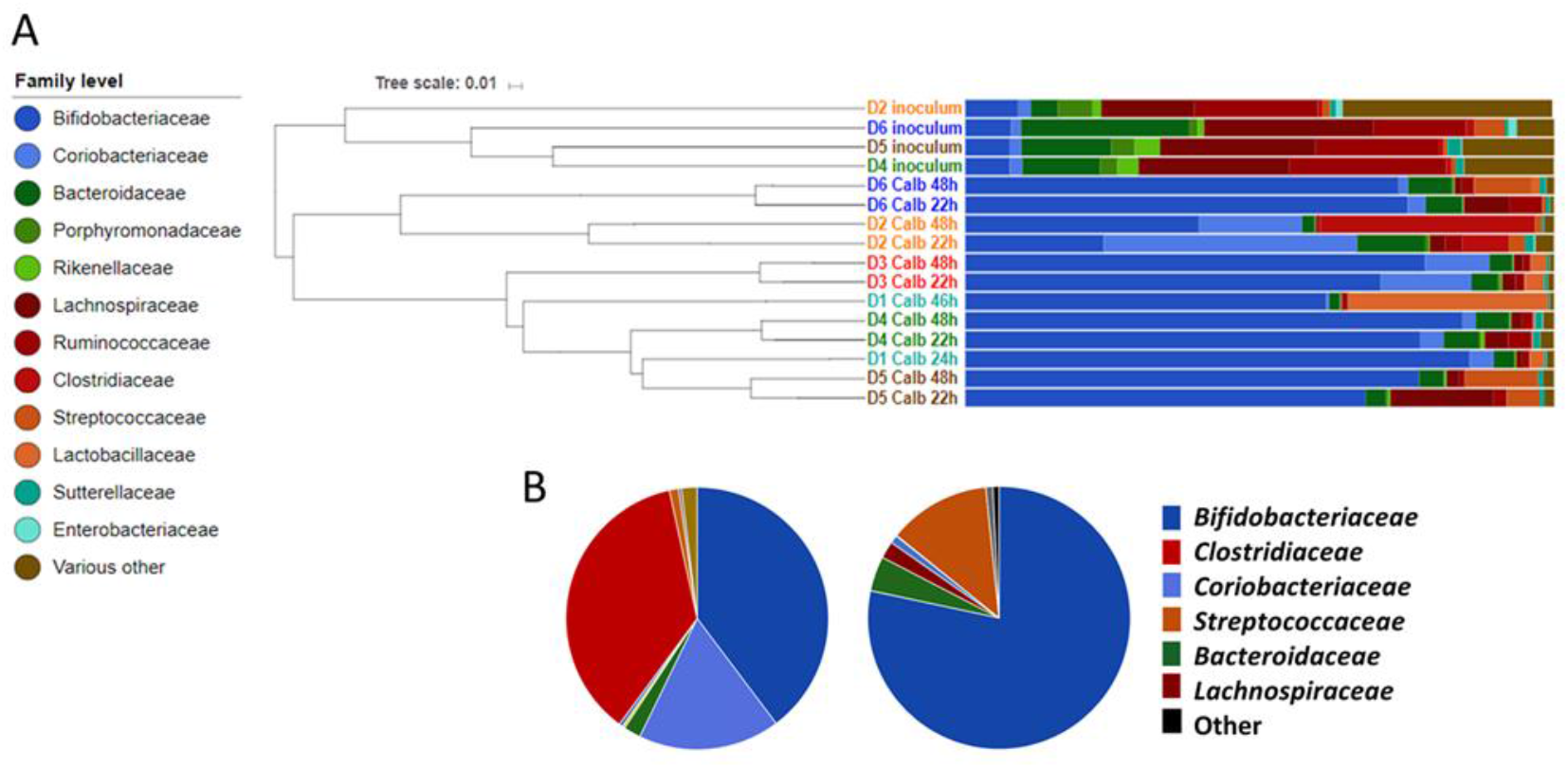
Bacterial community analysis of faecal samples and co-cultures with varying inhibitory activity against *C. albicans*. A) Bray–Curtis dendrogram of faecal inocula and subsequent co-cultures with *C. albicans*. B) Proportional family-level composition of Donor 2 (benign) and Donor 5 (antagonistic) faecal samples after 48 h co-culture with *C. albicans* in anaerobic, colon-mimicking, medium.

We classified the cultured faecal samples into different groups according to the observed impact on *C. albicans* growth in the batch co-culture. Specifically, Donor 5 was defined as ‘antagonistic’ as the faecal inoculum from this donor resulted in the strongest inhibitory effect, as were Donors 1, 3, 4, and 6 (all >85% *C. albicans* inhibition). The Donor 2 inoculum was classified as ‘benign’ since co-incubation had comparatively little effect on *C. albicans* survival *in vitro* (12% inhibition).

The non-parametric analysis of molecular variance (AMOVA) test implemented in the Mothur software package (Schloss *et al*. 2009) was first used to compare the bacterial compositions of the cultivated benign and antagonistic samples (D2 v D1, 3, 4, 5, 6) at days one and two combined revealed a statistically significant difference between the two groups (P=0.02).

We next used LEfSe (Segata *et al*. 2011) to identify taxa that were associated with either the antagonistic (D1, 3, 4, 5, 6) or benign status (D2). The analysis indicated that the *Bifidobacteriaceae* family (P=0.032), and more specifically, *Bifidobacterium adolescentis* (P=0.032) and *Bifidobacterium longum* derived OTUs (P=0.032) belonging to the Gram-positive *Actinobacteria* phylum correlated with samples exerting an antagonistic activity against *C. albicans* (**Figure 2B**; **Tables S4 and S5_Supplementary Data**). In contrast, the *Coriobacteriaceae* family (P=0.032) and the constituent species *Collinsella aerofaciens* (P=0.026) (hereon indicated as *Co. aerofaciens*), also belonging to the *Actinobacteria* phylum, together with *Clostridiaceae* (P=0.031) and *Clostridium neonatale* (P=0.026) from the *Firmicutes* phylum, correlated with the lack of antagonistic activity against *C. albicans* (**Figure 2B, Tables S4 and S5_Supplementary Data**).

### Culture supernatants of specific human gut isolates inhibit *C. albicans* growth under anaerobic conditions

Having correlated the presence of bifidobacteria in the cultivated faecal samples with antagonistic activity against *C. albicans* using the 16S rRNA gene-based analysis, we next attempted to verify this finding by testing a panel of 37 common and dominant gut bacterial strains for inhibition of *C. albicans* growth *in vitro*. The species selected for these tests were representative of the main phyla inhabiting the human gut (**Table S2_Supplementary Data**). The bacterial isolates of interest belonged to the phyla *Firmicutes* (nine strains belonging to the family *Lachnospiraceae*, four *Eubacteriaceae*, one *Peptostreptococcaceae*, three *Clostridiaceae*, six *Ruminococcaceae*, and one *Oscillospiraceae*), *Actinobacteria* (*B. adolescentis*, selected for analysis as this species was correlated with antagonist activity in co-culture with *C. albicans*, and two *Coriobacteriaceae*), *Bacteroidetes* (five *Bacteroidaceae*, one *Porphyromonadaceae*, and two *Prevotellaceae*) and one *Proteobacteria* (*Enterobacteriaceae*). A subset of the tested gut anaerobes was newly isolated for the purpose of this study from stool samples of healthy volunteers (see Materials and Methods section for details of isolation steps).

We reasoned that the inhibitory effects of gut microbes upon *C. albicans* might be mediated, at least in part, by secreted factors or metabolites. Therefore, in order to assess the putative *in vitro* inhibitory activity of the selected gut bacterial isolates, each species (**Figure 3**) was grown individually in M2GSC liquid medium overnight. Then, filter-sterilised culture supernatant was incubated with an overnight liquid culture of *C. albicans* under anaerobic conditions for 24 h. *C. albicans* biomass was assessed using optical density (OD_600_) measurements. The percentage growth of the fungus alone in fresh NGY medium, without exposure to bacterial supernatants, was set as 100% reference for each repeat run, and uninoculated M2GSC medium was used as a control.

**Figure 3:**
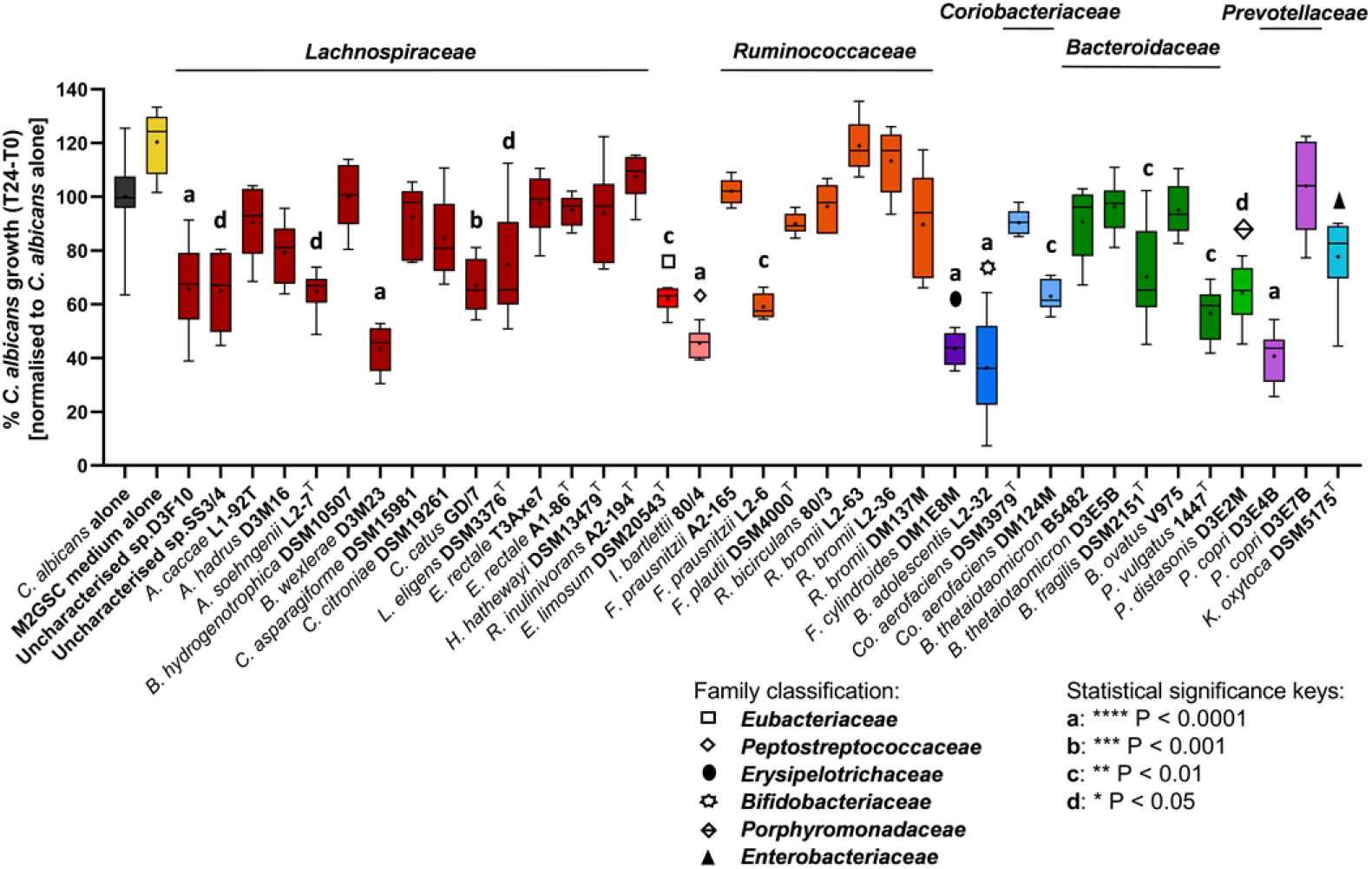
Impact of culture supernatants from individual human gut anaerobe strains on *C. albicans* growth under anaerobic conditions. The whisker boxplot represents percent *C. albicans* growth (T24–T0) when incubated with pure culture supernatants from human gut isolates. The growth of *C. albicans* alone in fresh NGY medium (black) was monitored via six technical replicates per test (total n=54). Strains are grouped by family and colour-coded: dark red for *Lachnospiraceae*; red for *Eubacteriaceae*; orange for *Ruminococcaceae*; purple for *Erysipelotrichia*; blue for *Bifidobacteriaceae*; light blue for *Coriobacteriaceae*; green for *Bacteroidaceae*; light green for *Porphyromonadaceae*; lilac for *Prevotellaceae*; and turquoise for *Enterobacteriaceae*. The cross represents the mean, while the central horizontal line shows the median of six technical replicates per strain (except for ‘Uncharacterised’ sp. D3F10, n=17; *Coprococcus catus* GD/7 and *Lachnospira eligens* DSM 3376^T^, n=12; *R. bromii* DM137M, n=11; *B. adolescentis* L2-32, n=24; *Bacteroides fragilis* DSM 2151^T^, n=11). The Kruskal–Wallis test revealed a highly significant difference between the effects of different supernatants (P<0.0001), and Dunn’s post-hoc identified multiple gut anaerobes whose culture supernatants significantly inhibited *C. albicans* growth compared to the *C. albicans*-only control, as indicated in the figure.

The experiments revealed that the different supernatants varied widely in their effect on *C. albicans* growth (**Figure 3**). Compared to controls, most of the isolates tested, including *Co. aerofaciens* DSM 3979^T^, which was correlated with benign status in the earlier sequence-based profiling analysis, did not inhibit *C. albicans* growth. Of note, however, *Co. aerofaciens* strain DM124M showed a mild inhibitory effect (P<0.01, **Figure 3**), suggesting that the activity observed may be strain specific. In contrast, the *Blautia wexlerae* D3M23, *Faecalitalea cylindroides* DM1E8M, *Prevotella copri* D3E4B, and *Intestinibacter bartlettii* 80/4 isolates showed more notable inhibitory effects (average inhibition in the range of 55–60%, P<0.0001) and *B. adolescentis* L2-32 was identified as the strongest antagonist among all of the strains tested (63.6% average inhibition, **Figure 3**, P<0.0001). This was consistent with the 16S rRNA gene-based analysis described above, which had associated bifidobacteria with inhibition of *C. albicans* in the co-culture experiments. Incubation with the bacterial growth medium alone (M2GSC) appeared to promote the growth of *C. albicans* slightly, although the effect was not statistically significant (**Figure 3**), likely due to the presence of glucose in the medium, which *C. albicans* can use for growth.

Because of the strong inhibitory impact displayed by the *B. adolescentis* strain L2-32 supernatant, combined with the previously identified correlation of this species with strong antagonism against *C. albicans* in the co-culture faecal incubation experiments described above, and the fact that this species is commonly detected in faeces from healthy adults (Matsuki *et al*. 2004), we next decided to focus on *Bifidobacterium* isolates and, in particular, on *B. adolescentis*, in more detail.

### Supernatants from specific *Bifidobacterium* strains inhibited *C. albicans* growth under anaerobic conditions

To investigate whether different species of bifidobacteria inhibited the growth of *C. albicans in vitro*, four different bifidobacterial species, including one *B. animalis* strain, four *B. adolescentis*, two *B. bifidum*, and six *B. longum* strains, all isolated from the faeces of healthy adults (**Table S2_Supplementary Data**), were screened for inhibition of *C. albicans* growth using the anaerobic assay described above. As *Co. aerofaciens* was correlated with benign effects on *C. albicans* in the faecal co-culture work, we also included supernatants from one strain of this species in these experiments for comparative purposes. The supernatants of all bifidobacteria species tested resulted in 20-80% *C. albicans* growth inhibition (relative to *C. albicans*-only growth in fresh NGY medium), except for *B. animalis* T1-817, which had no inhibitory activity (**Figure 4**). In agreement with the earlier experiments, supernatants from three out of four *B. adolescentis* strains (L2-32, L2-52, and L2-78) most strongly inhibited *C. albicans* growth (P<0.001; 68–78% fungal inhibition compared to the no supernatant controls) (**Figure 4**). In contrast, the type strain *B. adolescentis* DSM 20083^T^ did not show a strong inhibitory effect, further indicating that the inhibitory activities may be strain-specific. Supernatants from *B. bifidum* T2-126 and T2-106 cultures were also significantly antagonistic against *C. albicans* in the anaerobic assay (P<0.01 and P<0.001, with 42–49% fungal growth inhibition compared to the control, respectively). Finally, all representatives of the *B. longum* species tested showed a consistent, non-significant, mild inhibitory effect of approximately 20– 30% (**Figure** 4).

**Figure 4:**
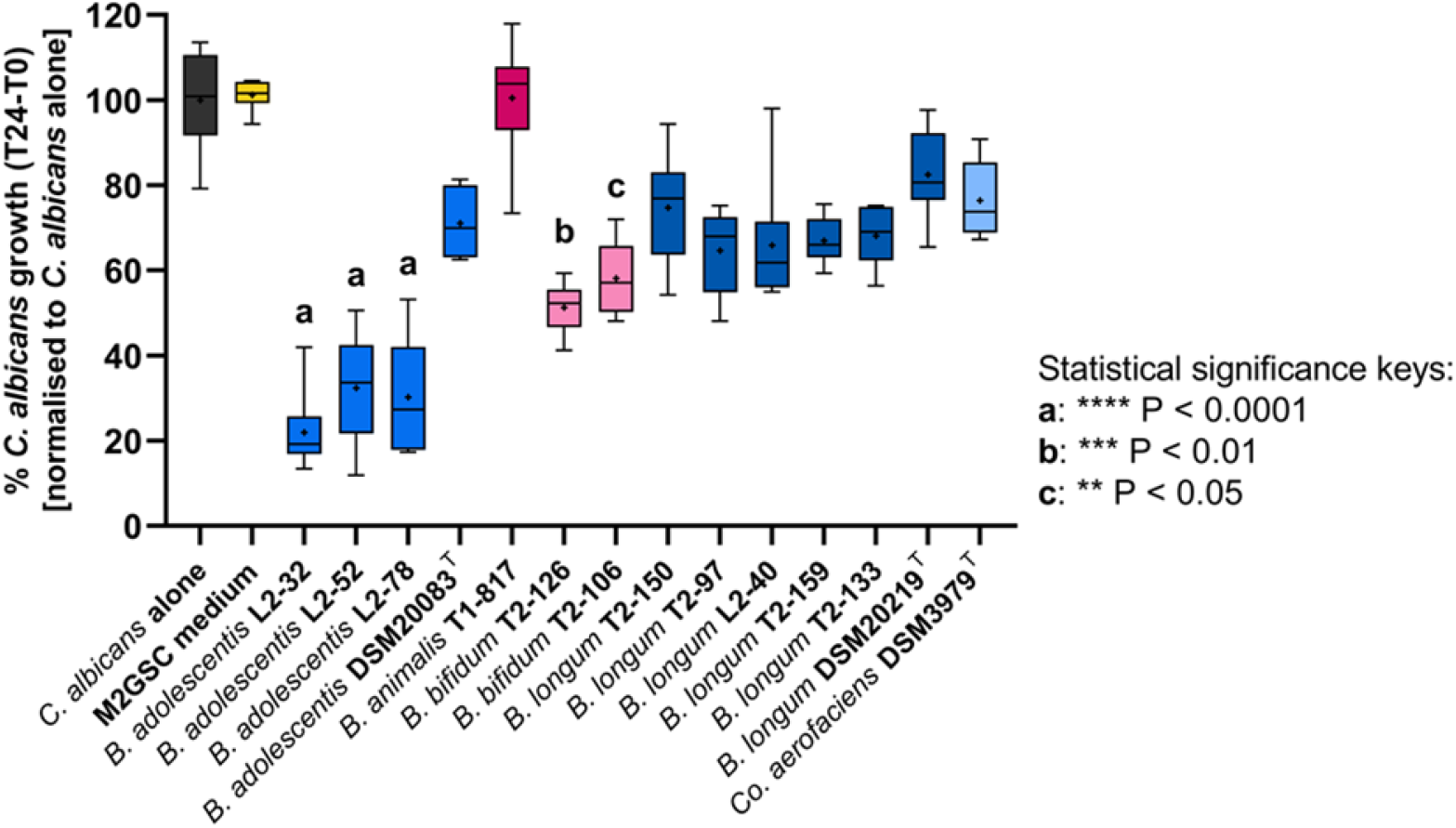
Impact of bifidobacterial and *Co. aerofaciens* culture supernatants on *C. albicans* growth under anaerobic conditions. The whisker boxplot represents % *C. albicans* growth (T24–T0) after incubation with culture supernatants from *Bifidobacterium* spp. or *Co. aerofaciens* strains isolated from healthy human donors. The crosses and central horizontal lines represent the mean and median, respectively, of six technical replicates per strain or for the *C. albicans*-only control (black). Strains are colour-coded by species. The Kruskal–Wallis test revealed a highly significant difference between samples (P<0.0001), and Dunn’s post-hoc test identified specific bifidobacterial isolates that exerted a significant inhibitory effect on *C. albicans* growth compared to the control.

### The inhibitory activity of bifidobacterial supernatants on *C. albicans* growth correlated with fermentation acid production and acidic pH

Having determined that culture supernatants from certain *Bifidobacterium* species from the human gut exert inhibitory activity against *C. albicans*, we next investigated the potential mechanisms underlying this phenomenon. As anticipated, quantification of the fermentation acids in the bifidobacterial supernatants used in the anaerobic assay revealed that the main organic acids produced by these strains were acetate, lactate, and formate (**Table S6_Supplementary Data**). *B. adolescentis* L2-32 produced the highest levels of the fermentation acids (38.1 mM acetate, 9.9 mM lactate, and 4.2 mM formate), followed by *B. adolescentis* L2-52 (20.67 mM acetate, 8.2 mM lactate, and 4.69 mM formate), and *B. adolescentis* L2-78 (31.21 mM acetate, 11.42 mM lactate, and 6.23 mM formate) (**Table S6_Supplementary Data**). The bifidobacterial strains producing the highest total concentrations of these fermentation acids therefore also displayed the strongest antagonistic activity against *C. albicans* (**Figure 5**). In contrast, we detected low concentrations of organic acids in non-inhibitory strain supernatants, such as those from *B. animalis* T1-817 and from the *B. longum* strains (**Table S6_Supplementary Data**), suggesting that the inhibitory capacity of certain human gut bifidobacteria might be associated with the release of primary metabolites into the supernatant.

**Figure 5:**
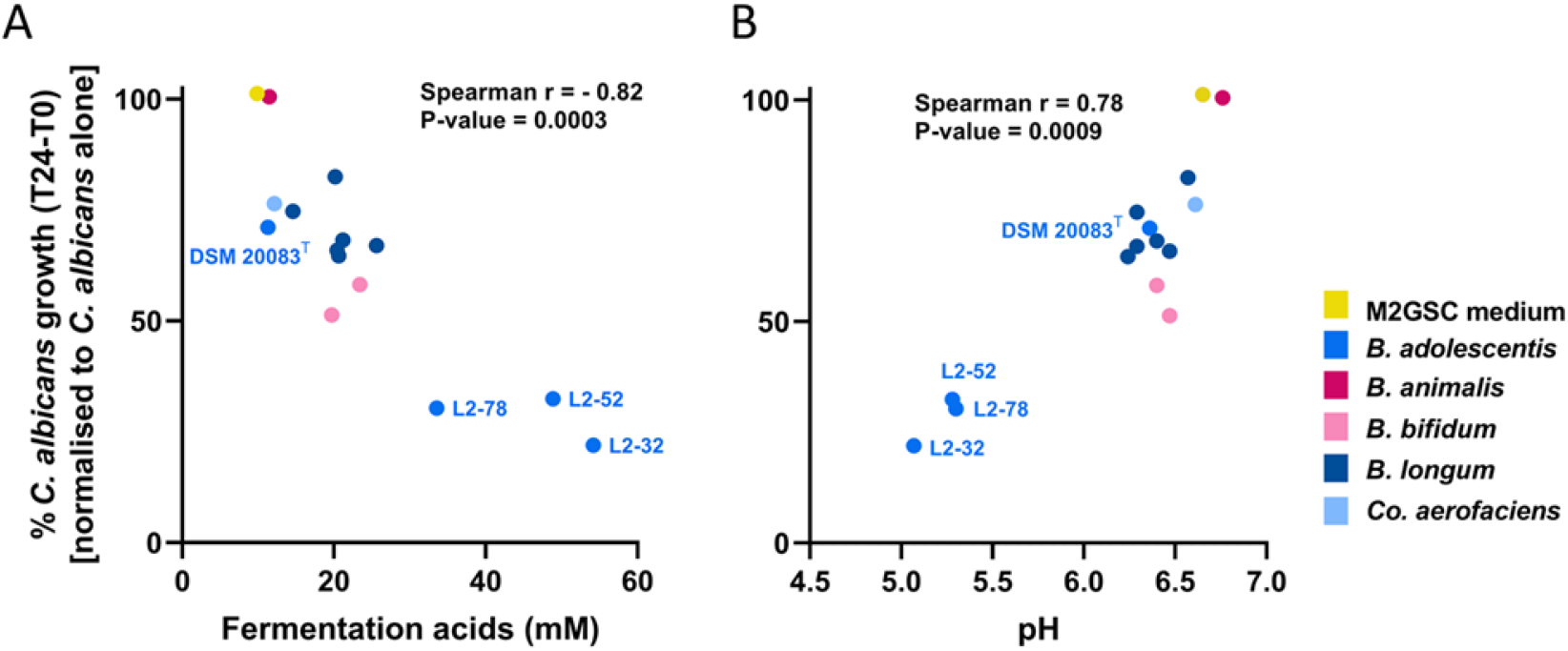
The inhibitory effect of *Bifidobacterium* and *Collinsella aerofaciens* isolates positively correlated with total concentration of fermentation acids and lower supernatant pH. Spearman correlation revealed that *C. albicans* inhibition was strongly associated with the fermentation acid concentration (A) and pH (B) of the bifidobacterial culture supernatants. Dots are colour-coded according to bacterial species, as per the key in the figure. P-values were corrected using the Benjamini, Krieger and Yekutieli false discovery rate approach.

To assess whether the inhibitory activity observed in the anaerobic assay was associated with the production of fermentation acids, we performed Spearman’s coefficient analysis by plotting the percent growth of *C. albicans* vs. the total amount of fermentation acids in the gut bacteria supernatants. We observed a strong positive correlation between total fermentation acid levels and fungal growth suppression (r=–0.82) (**Figure 5A**). Similarly, we noted a strong negative correlation between pH and *C. albicans* growth, with the lower pH correlating with reduced fungal growth (r=0.78) (**Figure 5B**).

We also calculated Spearman’s correlation coefficients for the main individual fermentation acids produced by the *Bifidobacterium* strains (**Table S6_Supplementary Data**). The analysis revealed that acetate, lactate, and formate concentrations were all significantly associated with *C. albicans* inhibition.

### Sensitivity of *C. albicans* to individual and combined fermentation acids, and pH extremes, under anaerobic growth conditions

We next tested the effect of individual and mixed fermentation acid solutions, at concentrations analogous to the previously observed highly inhibitory *B. adolescentis* supernatants (40–50 mM acetate, 10–15 mM lactate, and 10 mM formate), on *C. albicans* growth in the anaerobic assay. The fermentation acid mixture containing acetate, lactate, and formate significantly reduced *C. albicans* growth compared to the control over the incubation period (mean fungal inhibition of 38%, P<0.001; **Figure 6A**). Similarly, the individual fermentation acids showed a consistent suppressive effect on *C. albicans* growth (mean fungal inhibition of approximately 35% compared to controls), despite formate and lactate being added at lower concentrations than acetate (**Figure 6A**). This may be related to the fact that lactate and formate are stronger acids (pKa around 3.8) than acetate (pKa of 4.8). However, of note, the extent of inhibition exerted by the individual and mixed fermentation acid solutions was inferior to the impact on fungal growth displayed by *B. adolescentis* L2-32 supernatants in the same test (**Figure 6A**). This suggests the potential existence of additional inhibitory factors in the supernatant.

**Figure 6:**
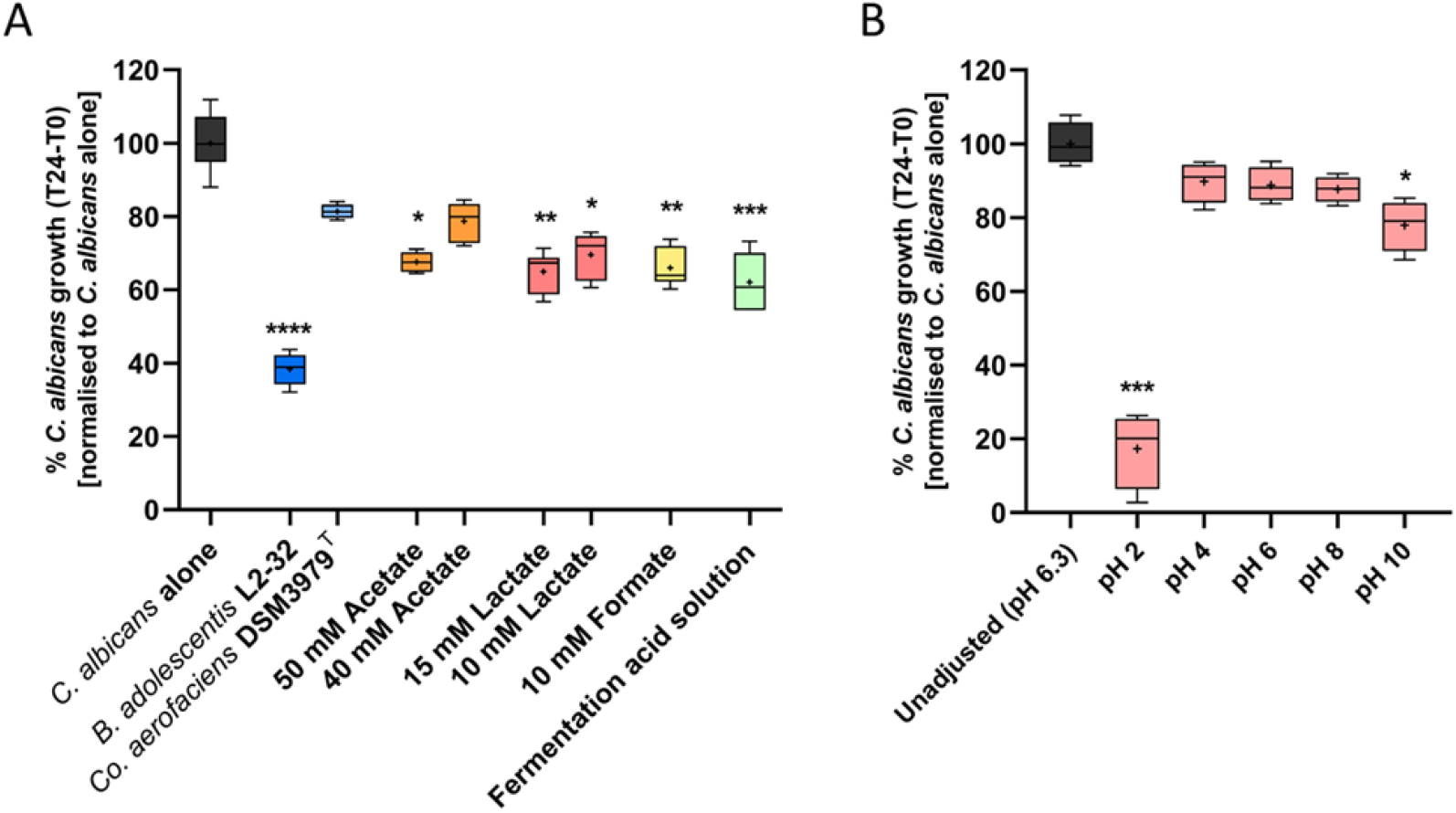
Impact of single and combined fermentation acids, as well as pH, on *C. albicans* growth under anaerobic conditions. A) Individual fermentation acids and a mixed acid solution at concentrations detected in the most inhibitory (*B. adolescentis* L2-32) supernatant (40 mM acetate, 10 mM lactate, and 10 mM formate) were tested for their impact on the growth of *C. albicans*. The whisker boxplot includes the mean and median of six technical replicates as crosses and horizontal lines, respectively. The Kruskal–Wallis test indicated strong differences between the observed values (P<0.0001); Dunn’s post hoc test revealed concentration-dependent inhibitory effects of the individual fermentation acids, with a particularly strong effect of 15 mM lactate and 10 mM formate, compared to the *C. albicans*-only control. B) Effect of pH on *C. albicans* growth, under anaerobic conditions. pH values were adjusted by modifying NGY culture medium before filter-sterilisation. The whisker boxplots show mean and median of four technical replicates. The Kruskal–Wallis test indicated significant differences between the observed values (P=0.0024); Dunn’s post hoc testing indicated significant differences in fungal growth between the medium with unadjusted pH (pH 6.3, black), and pH 2 and pH 10. Significance values: **** P<0.0001, *** P<0.001, ** P<0.01, * P<0.05.

We then assessed the sensitivity of *C. albicans* to pH, by incubating in NGY culture medium adjusted to pH values ranging from 2 to 10. In contrast to the fermentation acids-based tests, pH values within the normal range of those detected in the lower gastrointestinal tract seemed to have little impact on *C. albicans* growth when tested as the sole variable (**Figure 6B**). Indeed, fungal growth was only significantly decreased at extreme pH values, particularly at pH 2 (P<0.001) and at pH 10 (P<0.05), compared to the fungal growth in unadjusted NGY medium (**Figure 6B**). This indicated that the suppression of *C. albicans* growth observed in the presence of culture supernatants is not driven solely by pH.

### Inhibition of *C. albicans* by bifidobacterial supernatants was mediated via the combined effects of pH and SCFAs

To further uncover the mechanisms underpinning the inhibitory capacity of the *B. adolescentis* strains tested, we next set up an anaerobic assay to study the effect of the following individual stressors on *C. albicans* growth: pH alone, exposure to a mixed solution of fermentation acids (45 mM acetate, 15 mM lactate, and 10 mM formate, to mimic the concentrations determined in the most inhibitory (*B. adolescentis* L2-32) supernatant), and bacterial culture supernatants. To better understand the combinatorial role of fermentation acid concentration and pH, we conducted the tests at different controlled pH values, in the range from 4 to 7, adjusting either the medium, or the test solution/supernatant.

Consistent with the previous observations (**Figure 6**), *C. albicans* was highly resilient to the pH range tested under anaerobic conditions (**Figure 7**). Critically though, altering the pH significantly impacted the inhibitory activity of the tested supernatants, and the fermentation acids mix. In all cases, these treatments were most inhibitory at the lowest pH tested (pH 4), and progressively lost potency against *C. albicans* as the pH increased (**Figure 7**). This indicated that pH and fermentation acids combine to produce an inhibitory effect on *C. albicans*.

**Figure 7:**
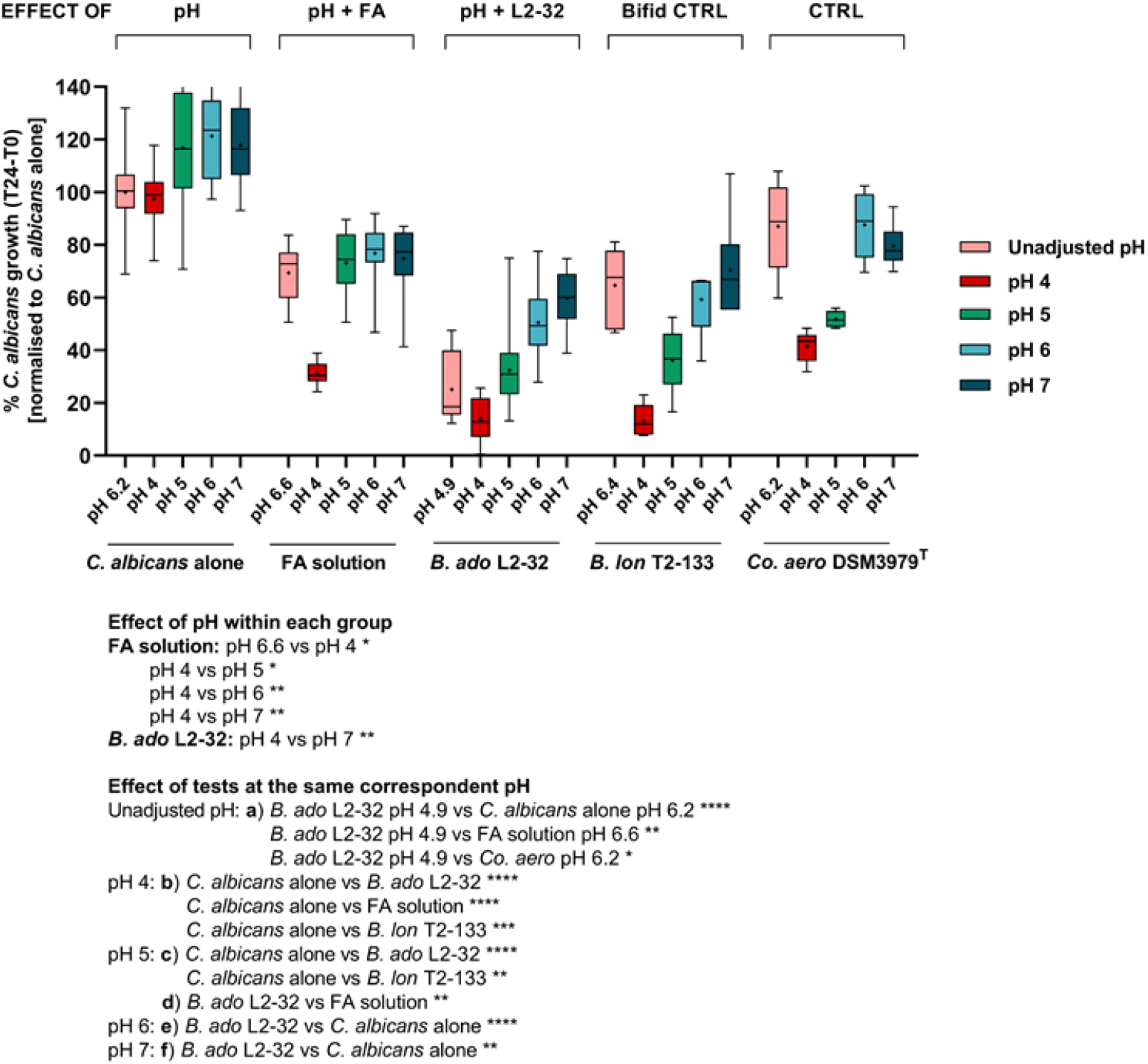
Cumulative impact of pH and fermentation acids (FA) on *C. albicans* growth under anaerobic conditions. The whisker boxplot shows *C. albicans* growth when tested at different controlled pH values, adjusting either the medium or the test solution/supernatant in the range from pH 4 to 7, under anaerobic conditions. Crosses and central horizontal lines represent the mean and median, respectively, of 12 technical replicates per test (n=32 for *C. albicans* alone at pH 6.2, n=18 for FA solution at all tested pH values, n=24 for both *B. adolescentis* L2-32 and *Co. aerofaciens* DSM 3979^T^ at all tested pH values). The Kruskal–Wallis test indicated highly significant differences between groups (P<0.0001); Dunn’s post-hoc test, comparing the observations against each other, indicated significant differences within each group at different pH values (colour-coded as per legend) and between groups at the same corresponding pH, indicated separately in the Figure legend as ‘effect of pH’ and ‘effect of tests at the same correspondent pH’, respectively. Significance: **** P<0.0001, *** P<0.001, ** P <0.01, * P<0.05. B. ado, *B. adolescentis*; B. lon, *B. longum*.

Of note, the antagonistic effect of the *B. adolescentis* L2-32 supernatant was significantly more pronounced than that of the SCFA solution at pH 5, as well as to that of a solution with an unadjusted pH value (**Figure 7**), again suggesting that the bacterial supernatant might contain additional inhibitory factors.

## DISCUSSION

*C. albicans* is a major clinical challenge because of high mortality in susceptible patients, emerging resistance against antifungal and sanitising agents, and the limited availability of additional therapeutic options (Pfaller *et al*. 2000). Alternative strategies to reduce carriage and dissemination of *C. albicans* in the gut should therefore be explored. The healthy intestinal microbiota is an appealing source of novel treatments, considering the well-established role it plays in protecting against systemic candidiasis, by hindering fungal expansion and pathogenic initiation in the gut (Kennedy and Volz 1985a, 1985b). However, the gut microbiota is extremely complex and it is currently largely unknown which components are most likely to be potent inhibitors of *C. albicans* in the gut. We demonstrate here that *B. adolescentis* culture supernatants exert strong inhibitory activity against *C. albicans* under anaerobic conditions *in vitro*, and identified an inhibitory effect of secreted bacterial fermentation acids and prevailing pH on *C. albicans* growth. These observations were in agreement with our DNA sequence-based analysis correlating the presence of *B. adolescentis* with the inhibition of *C. albicans* in mixed co-culture with faecal microbiota samples, under conditions mimicking the human colonic environment.

The *Bifidobacterium* genus is dominant in the colon of breast-fed infants (Yatsunenko *et al*. 2012; Khonsari *et al*. 2016) and it accounts for approximately 5% of the microbiota in adults, of which the species *B. adolescentis* is a prevalent representative (Reuter 1963). Importantly, *B. adolescentis* is also enriched following consumption of resistant starch (Ze *et al*. 2012), and produces high amounts of organic acids as a result of carbohydrate fermentation (**Table S6_Supplementary Data**). Despite the relatively low proportional abundance of this genus in the total microbiota in adults, it has potential health benefits for the host (Rossi *et al*. 2005; Fukuda *et al*. 2011; Rivière *et al*. 2014). Aside from fermentation acid production, bifidobacteria have also been demonstrated to induce the anti-inflammatory cascade (Lammers *et al*. 2003; Meng *et al*. 2016), and improve colonisation resistance against common food-borne pathogens such as *E. coli* O157:H7 and *Salmonella enterica* serovar Typhimurium (Makras and De Vuyst 2006; Fukuda *et al*. 2011; Ventura *et al*. 2016). In addition, *B. adolescentis* colonises the epithelial mucus layer and may therefore out-compete pathogens for adhesion sites on the gut epithelium (Tan *et al*. 2016; Ventura *et al*. 2016), potentially reducing the biofilm formation that can be an important virulence factor in *C. albicans* (Gulati and Nobile 2016).

Importantly, bifidobacteria were also recently predicted as major antagonists against *C. albicans* in an *in silico* model of inter-microbial interactions in the human gut (Mirhakkak *et al*. 2020). Bifidobacteria such as *B. adolescentis* may therefore be promising candidates for novel microbiota-based therapeutics aimed at enhancing colonisation resistance. Several clinical trials have reported some efficacy of probiotic supplementation of *Bifidobacterium* and *Lactobacillus* spp. in reducing *C. albicans* intestinal colonisation and preventing invasive fungal sepsis in infants following antibiotic treatment (Romeo *et al*. 2011; Roy *et al*. 2014). Furthermore, because *B. adolescentis* is a common member of the adult gut microbiota (present in up to 83% of healthy adults) (Matsuki *et al*. 1999; Junick and Blaut 2012) and responds to changes in the diet, the growth and metabolic activities of this species could potentially be modulated *in vivo* by prebiotic supplementation.

Aside from bifidobacteria, other gut bacterial taxa are also likely worthy of further study. For example, we also observed inhibitory effects against *C. albicans* by a number of other gut bacterial species (**Figure 3**). Wider screening of gut bacterial isolates is therefore highly likely to identify additional candidates with anti-*Candida* activity. In contrast, we also identified bacterial supernatants with little effect on *C. albicans* growth, such as those derived from *Flavonifractor plautii* and *Hungatella hathewayi*. This is consistent with reports that the relative abundances of these two bacterial species are correlated with *C. albicans* levels in faecal samples from cancer patients (Mirhakkak *et al*. 2020). Our results also highlight that different strains of the same gut bacterial species may have varying impacts on *C. albicans* growth (**Figure 3**). Better understanding of the mechanistic basis for some of the putative interactions, both beneficial and detrimental, between specific gut bacteria and *C. albicans* may help to prioritise candidates for development as novel therapeutics.

A key mechanistic result of the current study is demonstrating the combinatorial effect of fermentation acids and pH on the growth of *C. albicans*. Our findings are consistent with previous work indicating that the protonated form of weak acids freely permeate and accumulate inside microbial cells, causing dissipation of the proton motive force (Axe and Bailey 1995), triggering energetically expensive stress responses (Henriques, Quintas and Loureiro-Dias 1997), and perturbation of essential metabolic reactions (Cottier *et al*. 2015; Lourenço *et al*. 2018). Bacterial fermentation acids are therefore thought to play important roles in limiting *C. albicans* intestinal colonisation *in vivo* (Huang *et al*. 2011; Guinan *et al*. 2019), and a decrease in caecal SCFA concentrations following antibiotic treatment is associated with increased *C. albicans* load in the faeces in mouse models (Bohnhoff, Miller and Martin 1964; Guinan *et al*. 2019).

In agreement with our observations presented here, *C. albicans* was shown to be susceptible to formate (Mirhakkak *et al*. 2020) and acetate, at concentrations of over 30 mM, *in vitro*, and the effect is aggravated by microaerophilic conditions (Lourenço *et al*. 2018). Further, acetate inhibits hyphal morphogenesis of *C. albicans*, which is required for fungal translocation through the epithelial barrier (Guinan *et al*. 2019). In contrast, previous work has shown that lactate does not impair fungal growth at concentrations tested in our study, even at low pH values, and under aerophilic/microaerophilic conditions (Lourenço *et al*. 2018). Indeed, lactate is a potential energy source of *C. albicans* under hypoxic conditions, and is known to induce sustained fungal resistance to osmotic and cell wall stress, via cell wall remodelling (Ene *et al*. 2012b, 2012a, 2015). Nonetheless, substantial lactate release (up to approximately 110 mM), among other factors, is postulated to contribute to lactic acid bacteria-mediated colonisation resistance to *C. albicans* in the vaginal tract (Köhler, Assefa and Reid 2012; Zangl *et al*. 2020).

Importantly, the total fermentation acid and acetate concentrations that *C. albicans* cells were exposed to in this study are physiologically relevant for regions of the human gastrointestinal tract such as the proximal colon (Cummings and Macfarlane 1991). Indeed, total SCFA levels may reach up to 200 mM in the proximal colon (Cummings and Macfarlane 1991), suggesting that inhibition of *C. albicans* growth mediated by total fermentation acids may be greater than indicated by our study, and may represent a key mechanism of colonisation resistance to this opportunistic fungus. In contrast, the concentrations of formate and lactate detected here in the bifidobacterial culture supernatants appear to be slightly higher than those detected in human faecal samples, where they do not usually exceed 5–10 mM, as they are absorbed by the host or utilised by other bacteria (Hove, Norgard Andersen and Mortensen 1994; Duncan *et al*. 2007). Additionally, the finding that supernatants were often more inhibitory than defined fermentation acid mixtures (**Figures 6 and 7**) suggests that additional anti-fungal substances may be produced by some gut anaerobes. This may be a worthwhile avenue for further study.

## CONCLUSIONS

In this *in vitro* study we identified specific components of the human gut microbiota, *B. adolescentis* in particular, as antagonistic against *C. albicans*. Inhibitory activity was predominantly driven by the release of fermentation acids, and the subsequent drop in ambient pH. The potential for altering the gut microbiota composition, for example by consumption of probiotics such as *B. adolescentis*, or increasing *in vivo* SCFA concentrations by consumption of dietary fibres such as resistant starch, are worthy of further study to determine whether these can bolster colonisation resistance against *C. albicans* in the gut.

## Supporting information

All other supplemental files

## DECLARATIONS

### Competing interests

The authors have no conflicts of interest to declare.

### Funding

Initial studies were funded from a Wellcome Institutional Strategic Support Fund (ISSF) Seed Corn Award [105625/Z/14/Z]. Thereafter, the research was funded by the Scottish Government’s Rural and Environment Science and Analytical Services (RESAS) division.

### Authors’ contributions

AWW, SHD, AJPB, and MDL conceived of the research and designed the experiments. JM, GED, and AC carried out the batch co-culture experiment and analysed the resulting data. LR performed the rest of the experiments and analysed the data. KM and LR isolated novel gut bacterial strains that were used in these experiments. LR, SHD, and AWW wrote the manuscript. All authors read and approved the final manuscript.

## Acknowledgements

We thank Dr Donna M. MacCallum for critical reading of the manuscript, the Centre for Genome-Enabled Biology and Medicine at the University of Aberdeen for carrying out the 16S rRNA gene sequencing, and Donna Henderson for GC analysis of bacterial fermentation acids. The authors also acknowledge the support of the Maxwell computer cluster funded by the University of Aberdeen.

## Notes

### Competing Interest Statement

The authors have declared no competing interest.

