## Supplementary material for "Human gut bifidobacteria inhibit the growth of the opportunistic fungal pathogen *Candida albicans*": All other supplemental files

**Table S1**: Full sequencing results, including OTU table, phylum, family and genus level results, and alpha-diversity measures for each sample. See separate Excel spreadsheet

**Table S2**: List of gut anaerobe strains screened for inhibitory activity against *C. albicans*.

| **Family** | **Strain** | **Isolation laboratory/source** | **Reference** |
| --- | --- | --- | --- |
| *Lachnospiraceae* | Uncharacterised sp. D3F10 | Rowett Institute | This study |
| *Lachnospiraceae* | Uncharacterised sp. SS3/4 | Rowett Institute | Rowett Institute |
| *Lachnospiraceae* | *Anaerostipes caccae* L1-92^T^ | DSMZ | (Schwiertz *et al.* 2002) |
| *Lachnospiraceae* | *Anaerostipes hadrus* D3M16 | Rowett Institute | This study |
| *Lachnospiraceae* | *Anaerobutyricum soehngenii* L2/7 ^T^ | Rowett Institute | (Barcenilla *et al.* 2000; Shetty *et al.* 2018) |
| *Lachnospiraceae* | *Blautia hydrogenotrophica* DSM 10507^T^ | DSMZ | (Bernalier *et al.* 1996) |
| *Lachnospiraceae* | *Blautia wexlerae* D3M23 | Rowett Institute | This study |
| *Lachnospiraceae* | *Enterocloster asparagiformis* DSM 15981 | DSMZ | (Mohan *et al.* 2006) |
| *Lachnospiraceae* | *Enterocloster citroniae* DSM 19261 | DSMZ | (Warren *et al.* 2006) |
| *Lachnospiraceae* | *Coprococcus catus* GD/7 | Rowett Institute | (Duncan *et al.* 2009; Reichardt *et al.* 2014) |
| *Lachnospiraceae* | *Lachnospira eligens* DSM 3376^T^ | DSMZ | (Holdeman and Moore 1974) |
| *Lachnospiraceae* | *Eubacterum rectale* T3 Axe7 | Rowett Institute | Rowett Institute |
| *Lachnospiraceae* | *Eubacterum rectale* A1-86^T^ | Rowett Institute | (Barcenilla *et al.* 2000) |
| *Lachnospiraceae* | *Hungatella hathewayi* DSM 13479^T^ | DSMZ | (Steer *et al.* 2001) |
| *Lachnospiraceae* | *Roseburia inulinivorans* A2-194^T^ | Rowett Institute | (Barcenilla *et al.* 2000) |
| *Eubacteriaceae* | *Eubacterum limosum* DSM 20543^T^ | DSMZ | (Eggerth 1935) |
| *Peptostreptococcaceae* | *Intestinibacter bartlettii* 80/4 | Rowett Institute | (Duncan *et al.* 2009) |
| *Ruminococcaceae* | *Faecalibacterium prausnitzii* A2-165 | Rowett Institute | (Barcenilla *et al.* 2000),(Duncan *et al.* 2002) |
| *Ruminococcaceae* | *Faecalibacterium prausnitzii* L2/6 | Rowett Institute | (Barcenilla *et al.* 2000),(Duncan *et al.* 2002) |
| *Ruminococcaceae* | *Flavonifractor plautii* DSM 4000^T^ | DSMZ | (Seguin 1928) |
| *Ruminococcaceae* | *“Ruminococcus bicirculans”* 80/3 | Rowett Institute | (Walker *et al.* 2008) |
| *Ruminococcaceae* | *Ruminococcus bromii* L2-63 | Rowett Institute | (Barcenilla *et al.* 2000) |
| *Ruminococcaceae* | *Ruminococcus bromii* L2-36 | Rowett Institute | (Barcenilla *et al.* 2000) |
| *Ruminococcaceae* | *Ruminococcus bromii* DM137M | Rowett Institute | This study |
| *Erysipelotrichaceae* | *Faecalitalea cylindroides* DM1E8M | Rowett Institute | This study |
| *Bifidobacteriaceae* | *Bifidobacterium adolescentis* DSM 20083^T^ | DSMZ | (Reuter 1963) |
| *Bifidobacteriaceae* | *Bifidobacterium adolescentis* L2-32 | Rowett Institute | (Barcenilla *et al.* 2000) |
| *Bifidobacteriaceae* | *Bifidobacterium adolescentis* L2-52 | Rowett Institute | (Barcenilla *et al.* 2000) |
| *Bifidobacteriaceae* | *Bifidobacterium adolescentis* L2-78 | Rowett Institute | (Barcenilla *et al.* 2000) |
| *Bifidobacteriaceae* | *Bifidobacterium animalis* T1-817 | Rowett Institute | (Barcenilla *et al.* 2000) |
| *Bifidobacteriaceae* | *Bifidobacterium bifidum* T2-126 | Rowett Institute | (Barcenilla *et al.* 2000) |
| *Bifidobacteriaceae* | *Bifidobacterium bifidum* T2-106 | Rowett Institute | (Barcenilla *et al.* 2000) |
| *Bifidobacteriaceae* | *Bifidobacterium longum* T2-150 | Rowett Institute | (Barcenilla *et al.* 2000) |
| *Bifidobacteriaceae* | *Bifidobacterium longum* T2-97 | Rowett Institute | (Barcenilla *et al.* 2000) |
| *Bifidobacteriaceae* | *Bifidobacterium longum* L2-40 | Rowett Institute | (Barcenilla *et al.* 2000) |
| *Bifidobacteriaceae* | *Bifidobacterium longum* T2-159 | Rowett Institute | (Barcenilla *et al.* 2000) |
| *Bifidobacteriaceae* | *Bifidobacterium longum* T2-133 | Rowett Institute | (Barcenilla *et al.* 2000) |
| *Bifidobacteriaceae* | *Bifidobacterium longum* DSM 20219^T^ | DSMZ | (Reuter 1963) |
| *Coriobacteriaceae* | *Collinsella aerofaciens* DSM 3979^T^ | DSMZ | (Eggerth 1935) |
| *Coriobacteriaceae* | *Collinsella aerofaciens* DM124M | Rowett Institute | This study |
| *Bacteroidaceae* | *Bacteroides thetaiotaomicron* B5482 | A. Salyers (UIUC, Urbana, IL) | (Russell *et al.* 2013) |
| *Bacteroidaceae* | *Bacteroides thetaiotaomicron* D3E5B | Rowett Institute | This study |
| *Bacteroidaceae* | *Bacteroides fragilis* DSM 2151^T^ | DSMZ | (Veillon and Zuber 1898) |
| *Bacteroidaceae* | *Bacteroides ovatus* V975 | T.R Whitehead (USDA, Peoria, IL) | (Russell *et al.* 2013) |
| *Bacteroidaceae* | *Phocaeicola vulgatus* DSM 1447^T^ | DSMZ | (Eggerth and Gagnon 1933) emend. (Hahnke *et al.* 2016) |
| *Porphyromonadaceae* | *Parabacteroides distasonis* D3E2M | Rowett Institute | This study |
| *Prevotellaceae* | *Prevotella copri* D3E4B | Rowett Institute | This study |
| *Prevotellaceae* | *Prevotella copri* D3E7B | Rowett Institute | This study |
| *Enterobacteriaceae* | *Klebsiella oxytoca* DSM 5175^T^ | DSMZ | (Flugge 1886) |

DSMZ, Leibniz Institute, German Collection of Microorganisms and Cell Cultures GmbH

**Table S3**: Culturing conditions used to obtain novel isolates.

| **Family (Class)** | **Strain** | **Isolation medium** |
| --- | --- | --- |
| *Lachnospiraceae (Clostridia)* | Uncharacterised sp. D3F10 | FAA plus 5% B |
| *Lachnospiraceae (Clostridia)* | *Anaerostipes hadrus* D3M16 | M2GSC plus 0.5% H and 0.5% M |
| *Lachnospiraceae (Clostridia)* | *Blautia wexlerae* D3M23 | M2GSC plus 0.5% H and 0.5% M |
| *‎Ruminococcaceae (Clostridia)* | *Ruminococcus bromii* DM137M | M2GSC plus 0.5% H and 0.5% M |
| *Erysipelotrichaceae (Erysipelotrichia)* | *Faecalitalea cylindroides* DM1E8M | 0.2% AXOS-M2 enrichment, M2GSC |
| *Coriobacteriaceae (Actinobacteria)* | *Collinsella aerofaciens* DM124M | M2GSC plus 0.5% H and 0.5% M |
| *Bacteroidaceae (Bacteroidia)* | *Bacteroides thetaiotaomicron* D3E5B | 0.2% AXOS-M2 enrichment, BHI |
| *Porphyromonadaceae (Bacteroidia)* | *Parabacteroides distasonis* D3E2M | 0.2% AXOS-M2 enrichment, M2GSC |
| *Prevotellaceae (Bacteroidetes)* | *Prevotella copri* D3E4B | 0.2% AXOS-M2 enrichment, BHI |
| *Prevotellaceae (Bacteroidetes)* | *Prevotella copri* D3E7B | 0.2% AXOS-M2 enrichment, BHI |

Abbreviations: FAA, Fastidious Anaerobic Agar; B, defibrinated horse blood; H, haemin; M, menadione; AXOS, arabinoxylan oligosaccharides; BHI, brain heart infusion.

**Table S4**: Faecal bacterial families significantly associated with either antagonistic or benign effects on *C. albicans* growth in batch co-cultures using LEfSe.

| Phylotype | Class | LDA | P-value | Phylotype family designation |
| --- | --- | --- | --- | --- |
| 01 | Antagonistic | 5.34 | 0.032 | *Bifidobacteriaceae* |
| 02 | Benign | 5.13 | 0.032 | *Coriobacteriaceae* |
| 06 | Benign | 5.03 | 0.031 | *Clostridiaceae* |
| 18 | Benign | 3.37 | 0.005 | *Bacteroidales* unclassified |
| 30 | Benign | 3.71 | 0.025 | *Verrucomicrobiaceae* |
| 35 | Benign | 3.59 | 0.025 | *Bacteroidetes* unclassified |

Abbreviation: LDA, linear discriminant analysis, which estimates the effect size of each differentially abundant feature

**Table S5**: Faecal bacterial OTUs significantly associated with either antagonistic or benign effects on *C. albicans* growth in batch co-culture using LEfSe.

| OTU | Class | LDA | P-value | Representative OTU sequence  species designation |
| --- | --- | --- | --- | --- |
| Otu01 | Antagonistic | 5.17 | 0.032 | *Bifidobacterium adolescentis* |
| Otu04 | Antagonistic | 4.82 | 0.032 | *Bifidobacterium longum* |
| Otu05 | Benign | 5.09 | 0.026 | *Collinsella aerofaciens* |
| Otu06 | Benign | 5.06 | 0.026 | *Clostridium neonatale* |
| Otu27 | Benign | 2.71 | 0.049 | *Holdemanella biformis* |
| Otu35 | Benign | 3.96 | 0.005 | *Streptococcus anginosus* |
| Otu37 | Benign | 3.00 | 0.005 | *Bacteroidales* unclassified sp. |
| Otu44 | Antagonistic | 3.24 | 0.032 | *Bifidobacterium longum* |
| Otu51 | Antagonistic | 3.11 | 0.031 | *Bifidobacterium* unclassified sp. |
| Otu52 | Benign | 2.64 | 0.047 | *Blautia obeum* |
| Otu55 | Benign | 2.57 | 0.005 | *Odoribacter* unclassified sp. |
| Otu82 | Benign | 2.37 | 0.001 | *Ruminococcaceae* unclassified sp. |
| Otu84 | Benign | 2.49 | 0.001 | *Ruminococcaceae* unclassified sp. |
| Otu89 | Benign | 2.36 | 0.022 | *Phocaeicola dorei* |
| Otu98 | Benign | 2.12 | 0.005 | *Coprococcus* unclassified sp. |
| Otu103 | Benign | 2.68 | 0.011 | *Burkholderiales* unclassified sp. |
| Otu187 | Antagonistic | 2.17 | 0.030 | *Bacteroides uniformis* |
| Otu194 | Benign | 2.79 | 0.011 | *Erysipelatoclostridium ramosum* |
| Otu227 | Benign | 2.74 | 0.005 | *Bifidobacterium pseudocatenolatum* |
| Otu258 | Benign | 2.07 | 0.025 | *Ruthenibacterium lactatiformans* |
| Otu277 | Benign | 2.46 | 0.025 | *Collinsella aerofaciens* |
| Otu331 | Benign | 2.46 | 0.025 | *Clostridium neonatale* |
| Otu459 | Benign | 2.07 | 0.025 | *Bifidobacterium pseudocatenolatum* |

**Table S6**: Total and individual fermentation acid concentrations in the culture supernatants of bifidobacterial strains, together with corresponding pH and % *C. albicans* growth (T24–T0; n=6 technical replicates).

| Strain | % *C. albicans* growth | Acetate (mM) | Lactate (mM) | Formate (mM) | Total fermentation acids | pH |
| --- | --- | --- | --- | --- | --- | --- |
| *B. adolescentis* L2-32 | 22.00 | 37.17 | 8.7 | 8.34 | 54.21 | 5.07 |
| *B. adolescentis* L2-52 | 32.43 | 20.67 | 8.2 | 4.69 | 33.56 | 5.3 |
| *B. adolescentis* L2-78 | 30.26 | 31.21 | 11.42 | 6.23 | 48.86 | 5.28 |
| *B. adolescentis* DSM 20083^T^ | 71.13 | 6.05 | 3.1 | 2.16 | 11.3 | 6.36 |
| *B. animalis* T1-817 | 100.48 | 7.64 | 0 | 3.81 | 11.45 | 6.76 |
| *B. bifidum* T2-126 | 51.30 | 13.84 | 2.77 | 3.1 | 19.71 | 6.47 |
| *B. bifidum* T2-106 | 58.17 | 16.22 | 4.07 | 3.12 | 23.41 | 6.4 |
| *B. longum* T2-150 | 74.74 | 13.83 | 4.12 | 3.24 | 21.19 | 6.4 |
| *B. longum* T2-97 | 64.68 | 17.62 | 5.45 | 2.53 | 25.6 | 6.29 |
| *B. longum* L2-40 | 65.94 | 14.62 | 4.32 | 1.71 | 20.64 | 6.24 |
| *B. longum* T2-159 | 67.03 | 14.02 | 4.59 | 1.55 | 20.17 | 6.57 |
| *B. longum* T2-133 | 68.17 | 12.95 | 3.87 | 3.58 | 20.41 | 6.47 |
| *B. longum* DSM 20219^T^ | 82.52 | 8.74 | 4.22 | 1.64 | 14.6 | 6.29 |
| *Co. aerofaciens* DSM 3979^T^ | 76.43 | 0.46 | 6.52 | 5.19 | 12.16 | 6.61 |
| M2GSC medium | 101.21 | 9.87 | 0 | 0.72 | 10.59 | 6.65 |
| Spearman coefficient | | -0.87 | -0.62 | -0.46 | -0.82 | 0.78 |
| Corrected P-value (two-tailed) | | 0.0001 | 0.0043 | 0.0175 | 0.0002 | 0.0003 |
